# mPPases create a conserved anionic membrane fingerprint as identified via multi-scale simulations

**DOI:** 10.1101/2022.03.08.483421

**Authors:** Alexandra O. M. Holmes, Adrian Goldman, Antreas C. Kalli

## Abstract

Membrane-integral pyrophosphatases (mPPases) are membrane-bound enzymes responsible for hydrolysing inorganic pyrophosphate and translocating a cation across the membrane. Their function is essential for the infectivity of clinically relevant protozoan parasites and plant maturation. Recent developments have indicated that their mechanism is more complicated than previously thought and that the membrane environment may be important for their function. In this work, we use multiscale molecular dynamics simulations to demonstrate for the first time that mPPases form specific anionic lipid interactions at 4 sites at the distal and interfacial regions of the protein. These interactions are conserved in simulations of the mPPases from *Thermotoga maritima, Vigna radiata* and *Clostridium leptum* and characterised by interactions with positive residues on helices 1, 2, 3 and 4 for the distal site, or 9, 10, 13 and 14 for the interfacial site. Due to the importance of these helices in protein stability and function, these lipid interactions may play a crucial role in the mPPase mechanism and enable future structural and functional studies.

**Author summary:** In this work we have been able to demonstrate conservation of lipid-interaction sites on proteins from distinct species that deviated from their evolutionary common ancestors a long time ago, as in the case of the membrane-integral pyrophosphatases from a thermophilic bacteria species and a plant species studied here. This retention of a common lipid interaction profile or “fingerprint” and our ability to predict this for other proteins in this family may indicate that they are more integral to protein function than previously thought. By identifying lipid interactions that may act to stabilise the protein structure, these properties could be exploited to gain protein structures, and the interfacial site’s potential involvement in inter-subunit communication may be useful for further investigation of the catalytic cycle of this clinically relevant membrane protein family.

## Introduction

Membrane integral pyrophosphatases (mPPases) are a family of membrane proteins responsible for coupling the hydrolysis of the pyrophosphate (PP_i_) phosphoanhydride bond to the pumping of a cation across the membrane (Holmes et al., 2019). This allows mPPases to both remove excess PP_i_ from the cytoplasm, and to generate a membrane potential. mPPases are found in all kingdoms of life, excluding fungi and multicellular animals (Kajander et al., 2013). Due to this, they are validated selectively toxic drug targets against a variety of protozoan and bacterial pathogens (Rodrigues et al., 2000; Lemercier et al., 2002; Yoon et al., 2013; Liu et al., 2014) and reducing the function of mPPases *via* novel inhibitors may play a role in combatting these infectious pathogens.

Crystal structures revealed that mPPases exist as homodimers, where each subunit is composed of 16 transmembrane helices (TMH) (Lin et al., 2012; Kellosalo et al., 2012; Li et al., 2016; Vidilaseris et al., 2019; Tsai et al., 2019). A single subunit is formed by two concentric rings of TMH: the inner ring (TMH 5, 6, 11, 12, 15 and 16) makes up the 4 catalytic regions: the catalytic centre, the coupling funnel, the ionic gate and the exit channel, while the outer ring (TMH 1, 2, 3, 4, 7, 8, 9, 10, 13, 14) forms the subunit-subunit interface and the membrane-facing surface of the protein. Despite their common structure, seven different mPPase subfamilies have been functionally characterised (Luoto et al., 2011; Tsai et al., 2014; Luoto et al., 2015). In short, mPPase catalytic activity is either K^+^ dependent or K^+^ independent and the pumping specificity is either H^+^ only, Na^+^ only, dual H^+^/Na^+^ or H^+^ regulated by Na^+^. Of these subfamilies, only the 3D structures of the K^+^-dependent H^+^-PPase from *Vigna radiata* (*Vr*-PPase) (Lin et al., 2012), and the K^+^-dependent Na^+^-PPase from *Thermotoga maritima* (*Tm*-PPase) (Kellosalo et al., 2012; Li et al., 2016; Vidilaseris et al., 2019) have been resolved to high resolution.

These 3D structures have facilitated understanding of the mechanism by which mPPases perform hydrolysis and ion pumping. However, there is conflict surrounding the order of these two events (Baykov, 2020; Anashkin et al., 2021) and more recently, data have been published indicating that the mechanism is more complicated than previously thought (Artukka et al., 2018; Vidilaseris et al., 2019). There is also conflict about how the mPPase subunits operate with one another, which may be explained by the environment of the protein: lipidated (Artukka et al., 2018) or detergent-solubilised (Vidilaseris et al., 2019), and so could indicate protein-lipid interactions (Holmes et al., 2019). Despite evidence that the lipid environment may have a role in regulating the function of mPPases, the interaction of mPPases with their lipid environment is still unknown.

In recent years, molecular dynamics (MD) simulations have played a role in uncovering and studying membrane protein-lipid interactions (Corradi et al., 2019). These interactions can have multiple effects on the protein of interest, for example modulating stability (Habeck et al., 2015), assisting conformational changes (Sweadner, 2017), oligomerisation (Gupta et al., 2017; Pyle et al., 2018) and largescale protein organisation (Kalli and Reithmeier, 2018). MD simulations represents a robust way to identify putative lipid binding sites which can be refined through further simulations or in tandem with other methods (Corradi et al., 2019).

In this study, we examined the interactions and dynamics of three mPPases structures in model lipid bilayers *via* multi-scale MD simulations. Our results suggest that mPPases form an anionic annulus in the membrane and possess specific anionic lipid binding sites at the dimer interface and the distal regions of the protein. These protein-lipid interactions are conserved between mPPases from different species with differing pumping specificities that may suggest that these are a general property of mPPases.

## Results

### *Tm*-PPase forms an anionic fingerprint in the membrane

We first performed simulations with the 3D structure of the *Tm*-PPase due to the extensive structural characterisation of the protein over the last decade (Kellosalo et al., 2012; Li et al., 2016; Vidilaseris et al., 2019). The *Tm*-PPase structure was inserted into bilayers containing POPE and POPA, POPG or POPS molecules (Table 1) (**Fig. 1**). Little is known about the specific compositions of the *T. maritima* native lipid bilayers (Völkl et al., 1993; Guan et al., 2013; Sohlenkamp and Geiger, 2015) and so palmitoyl-oleoyl phospholipids were considered an appropriate proxy. Following 5 μs of simulation, lipid contact analysis revealed that the anionic lipids POPA, POPG and POPS interacted preferentially with *Tm*-PPase in the cytoplasmic leaflet compared to the zwitterionic POPE lipids (**Fig. 2**). Titration of the concentration of each anionic lipid, (40%, 20% and 10% of the overall lipid concentration) demonstrated that this annulus was retained in all concentrations with (~20 lipids) interacting with the protein in each concentration. Therefore, the 20% anionic lipid bilayers were chosen for further experiments, so selective interactions could be distinguished.

**Fig. 1.**
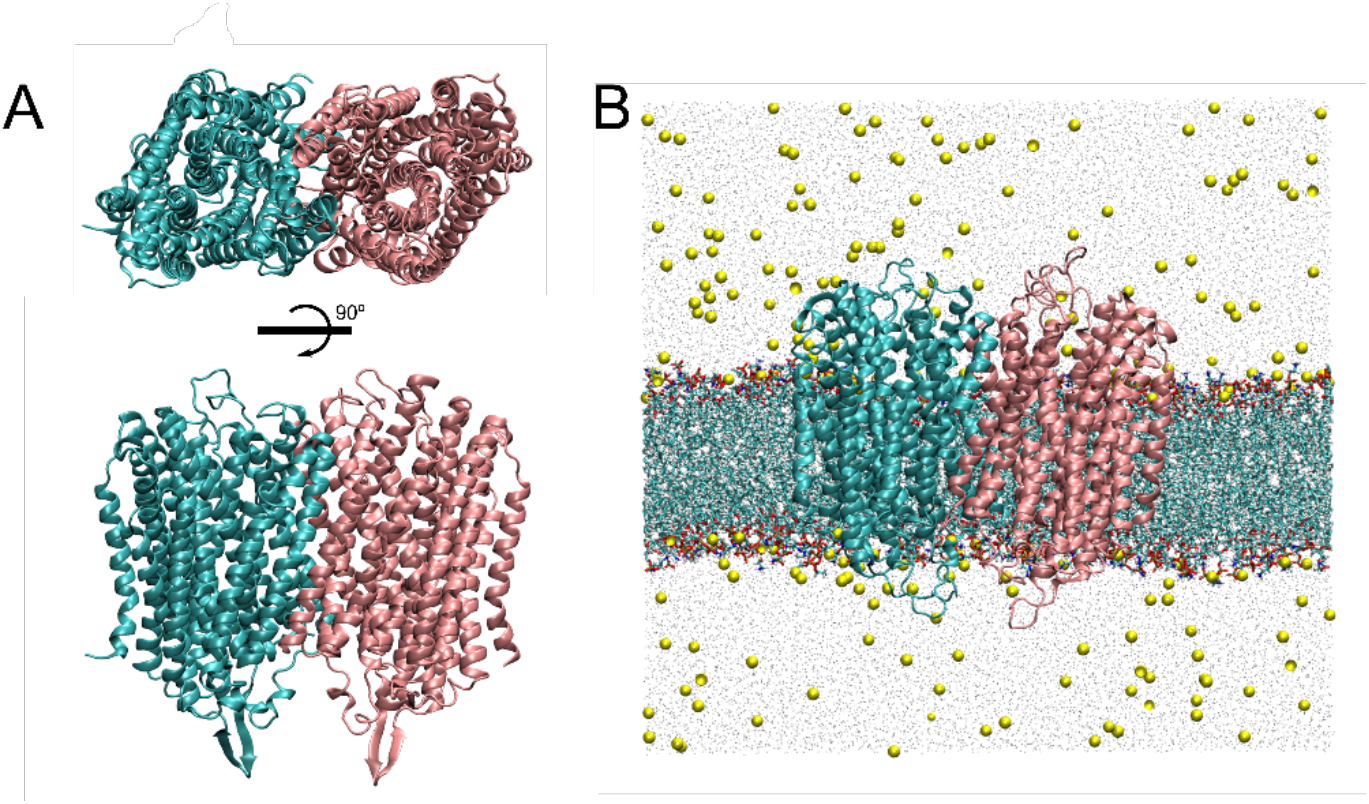
The structure of Tm-PPase and simulation box set up. A) the Tm-PPase resting state crystal structure (PDB: 4AV3) viewed from above to demonstrate the concentric circles of TMH containing the active site and ion translocation channel of each subunit (one cyan and one pink). B) The atomistic simulation box where the protein subunits are coloured as before and shown in cartoon, the POPE (80%) and POPA (20%) bilayer is shown as lines representation and the water particles are grey points, and the ions (NaCl 150 mM) are shown as yellow VDW particles.

**Fig. 2.**
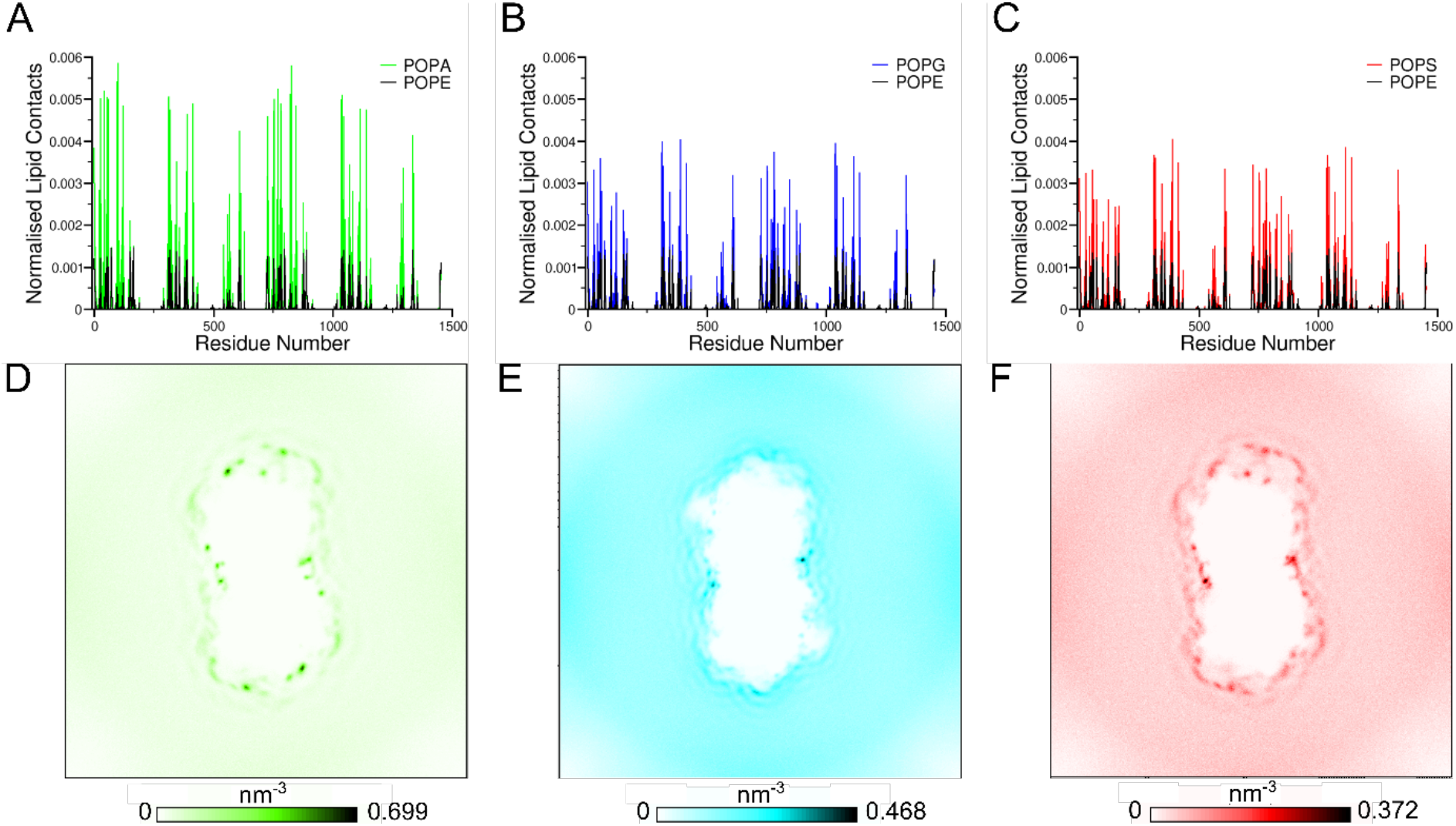
*Tm*-PPase interacts preferentially with anionic lipids at four distinct sites. Normalised number of contacts between the lipids and Tm-PPase in the simulations in which the model bilayers consist of 80% POPE and 20% A) POPA, B) POPG or C) POPS. The lipid contacts were normalised by dividing the number of contacts of each residue with the length of simulation and the number of lipid species. Average density of the phosphate particles of the anionic lipids D) POPA, E) POPG and F) POPS around the protein in the same simulations as in panels A-C.

Analysis of the interactions of each residue of the *Tm*-PPase with lipids revealed four symmetrical anionic lipid binding sites at the dimer interface and the “distal regions” of the protein (**Fig. 2**). Nine positive lysine and arginine residues (R^1.60^, K^1.61^, R^2.38^, K^3.59^, R^3.63^, K^4.40^, R^4.48^, K^8.40^ and K^8.42^) (note that the Ballesteros and Weinstein numbering system is used (Ballesteros and Weinstein, 1995; Tsai et al., 2014)) formed each of the distal interaction sites. The residues involved in the dimer interface sites were four positive lysine residues (K^9.70^, K^10.49^, K^13.52^ and K^14.45^), two of which were located in one subunit and the other two on the other subunit.

Additionally, POPA interacted more frequently with Tm-PPase compared to POPG and POPS at 20% anionic lipid bilayer content (**Fig. 2**). To examine further the more frequent binding of POPA lipids we have also performed a simulation in which the bilayer contained an equal anionic lipid mix (10% each of POPA, POPG and POPS). In this simulation, POPA interacted up to 43-fold or 84-fold more than POPG or POPS, respectively, augmenting our previous observation that POPA interact more with *Tm*-PPase compared to other anionic lipids.

### *Vr*-PPase in its native bilayer forms a similar membrane fingerprint

To examine whether the anionic lipid fingerprint identified above was *Tm*-PPase specific or if it also occurs in mPPases in other species and in other membranes, we also performed simulation using the crystal structure of the *Vr*-PPase. Like *Tm*-PPase, *Vr*-PPase is structurally well-characterised but mesophilic rather than thermophilic.

As we have more information about the *V. radiata* tonoplast membrane, simulations of *Vr*-PPase were performed in bilayers resembling this membrane (29% cholesterol, 25% POPC, 17% POPE, 17% ceramide hexoside, 6% PIP_2_, 3% POPG, 2% POPS and 1% POPA) (Yoshida and Uemura, 1986). The four anionic interaction sites seen with *Tm*-PPase were present at the interfacial and distal regions of the *Vr*-PPase protein (**Fig. 3**), where lysine and arginine residues composed the distal site (K^1.60^, K^1.62^, K^1.67^, K^2.50^, K^3.59^, R^4.33^, K^4.37^, R^4.44^, K^8.41^, K^8.52^). However, unlike *Tm*-PPase, the distribution of residues forming the interfacial site was uneven, as one from one subunit and three from the other contributed to forming interactions with the anionic lipids (K^10.49^, K^13.48^, K^14.40^, K^14.48^ for *Vr*-PPase). However, the arrangement of these four was highly similar to that seen in the *Tm*-PPase systems.

**Fig. 3.**
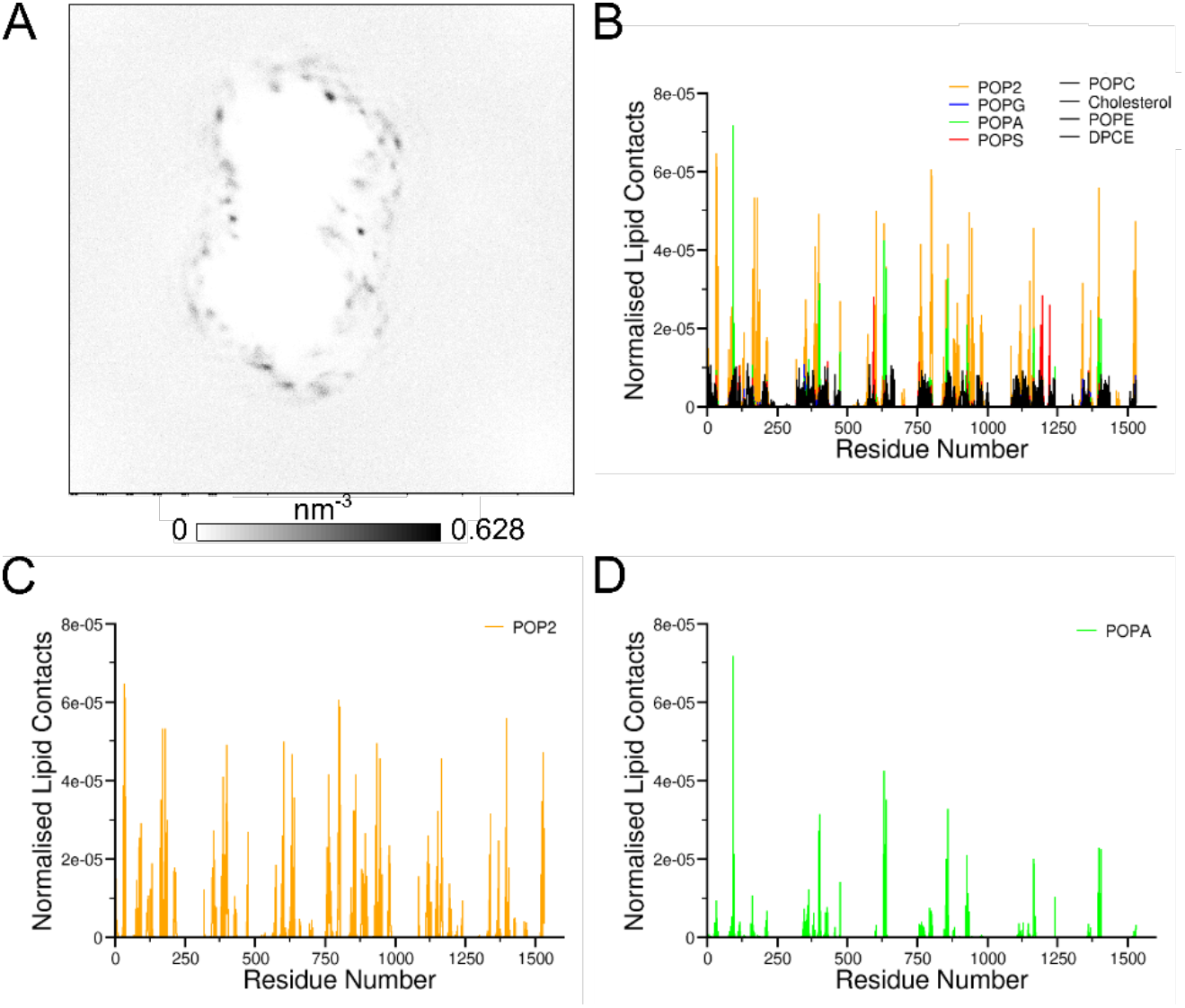
*Vr*-PPase in its native bilayer forms an anionic membrane fingerprint. A) Average density of the phosphate particles of the anionic lipids (PIP_2_, POPG, POPS and POPA). B-D) Normalized number of contacts between the lipids in the simulation of the *Vr*-PPase with the native-like bilayer.

These simulations contained additional anionic lipids compared to the systems with *Tm*-PPase, allowing us to study the preference of *Vr*-PPase for anionic lipids. All the anionic lipids formed interactions with the interfacial and distal interaction sites, but our analysis suggests that interactions of *Vr*-PPase with PIP_2_ were favoured over the neutral and other anionic lipids. Of the palmitoyl-oleoyl phospholipids, POPA was the favoured lipid over both POPG and POPS, despite its low representation in the membrane (1%).

### Homology-modelled crystallography target *Cp*-PPase retains this fingerprint

As the anionic lipid fingerprint was seen in two structurally characterised proteins and the indication from the literature that lipid binding could be of interest for functional and stabilising studies, we have also modelled the Na^+^/H^+^-PPase from *C. leptum* (*Cp*-PPase; see Methods for details). *Clostridium* species are predicted to contain POPG, POPE, POPS and cardiolipin, in addition to diradylglycerols. For this reason, we simulated this structure in the same bilayers as *Tm*-PPases. This also makes comparison of our results easier. *Cp*-PPase we simulated with 20% POPA, POPG or POPS or an equal anionic lipid mix (10% each). *Cp*-PPase is also a different type of mPPase; this will allow us to examine whether the anion interaction identified for two different mPPases are retained to other families.

As in the other systems, *Cp*-PPase formed an anionic lipid fingerprint in the membrane, where specific interactions formed between the lipids and positively charged residues at the distal site (R^1.56^, K^1.60^, K^2.46^, R^2.47^, K^3.59^, R^7.62^, K^7.64^) and at the interfacial site (K^9.69^, K^9.73^, R^13.52^, R^14.48^). As with *Tm*-PPase, the interfacial site was made up of two residues from each protein subunit. Again, *Cp*-PPase displayed preference for POPA over the anionic lipids POPG or POPS in the mixed simulations. This augments our previous observation that binding of anionic lipids to two sites may be a general property for all PPases.

### *In silico* mutations disrupt the protein-lipid interactions

To examine whether there was any synergy in the binding of lipids in the lipid binding sites identified above, we performed in-silico mutagenesis of the four positively charged residues identified above in the interfacial site to alanine. Analysis of the lipid density following identical simulation to the wild-type (WT) proteins showed that when a single or double site mutation (DSM) was made, the anionic lipid binding was lost, but binding at the intact site remained. In single site mutants, this was also seen, but only at the mutated site, indicating that our simulations do not show any cooperativity between the binding sites. This confirmed that the anionic lipid binding at these sites was due to specific interactions with the protein, rather than a stochastic effect, and that the binding is independent at each site. To assess whether this was due to the properties of the protein surface in these areas, electrostatic analysis of the proteins was performed and revealed that positively charged regions matched the location of the anionic binding at the interfacial and distal regions of the protein (**Fig. 4**). The electrostatic profiles were also similar between the three mPPases we have used in this study.

**Fig. 4.**
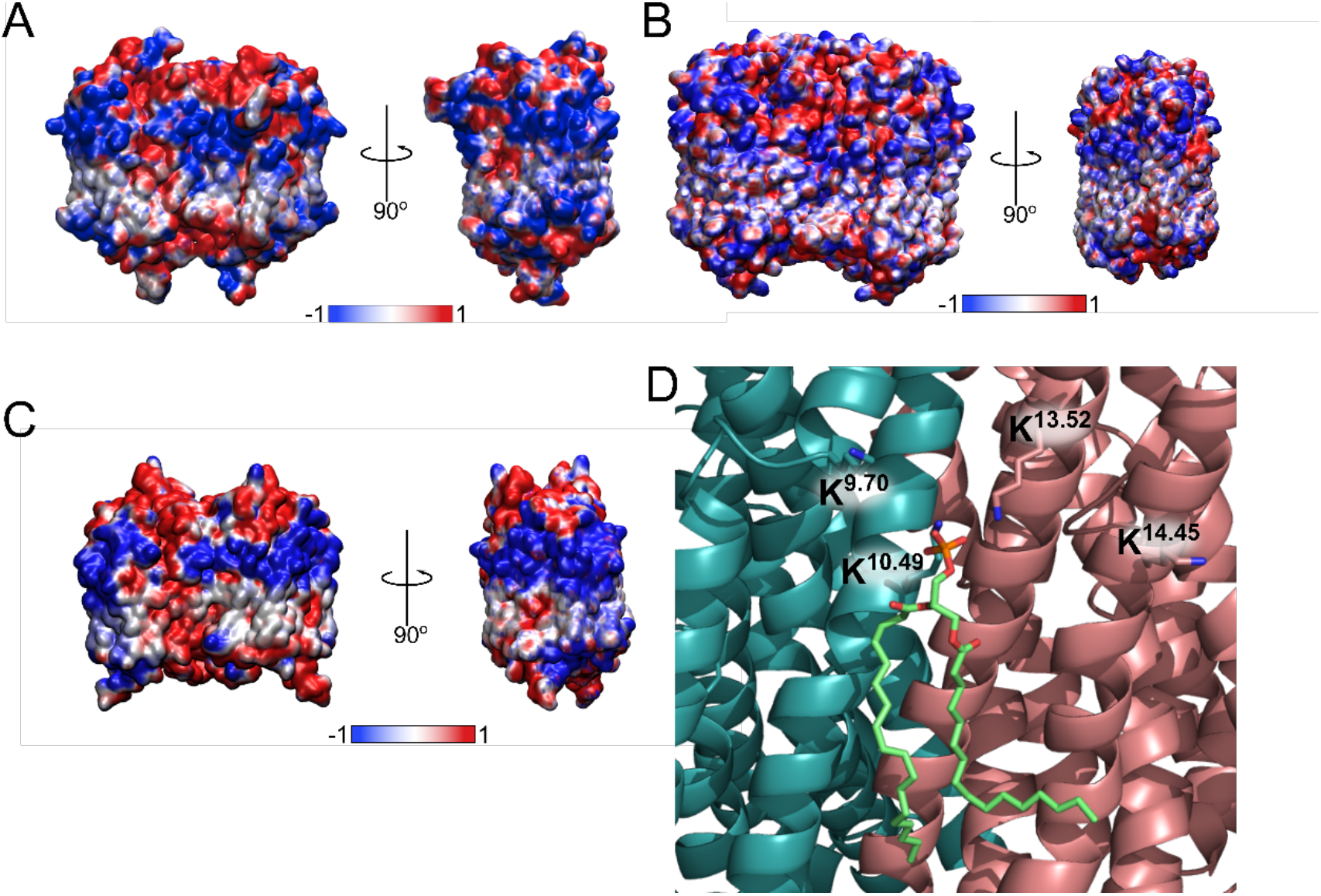
Electrostatic profiles of mPPases reveal positive charges at the anionic lipid binding sites. A-C) The electrostatic profiles of the *Tm*-PPase, *Vr*-PPase and *Cp*-PPase structures as calculated by APBS, respectively. D) Atomistic detail of the *Tm*-PPase interfacial anionic binding site with POPA (green sticks) bound and the interacting lysines shown as sticks. QuickSurf is used for the electrostatic profile representation and coloured by electrostatic profile (blue = positive, red = negative), cartoon representation is used for binding site detail.

### Refinement of the lipid interactions *via* atomistic simulations

The effect of lipid binding on the dimeric arrangement or stability of the protein could not be assessed through these CG simulations, as the elastic network between the protein monomers maintained the protein-protein interface and conformational state. Therefore, to refine the protein-lipid interactions at these sites and identify any effect the lipid binding had on protein structure and dynamics, representative final frames of the WT or DSM *Tm*-PPase, *Vr*-PPase and *Cp*-PPase systems were converted to an AT representation (see Methods).

The association of the anionic lipids at the WT protein interface was maintained throughout the 100 ns simulations, and interactions with all the proposed lysine and arginine residues at the mPPase protein interfaces were retained. Analysis of the simulations revealed interacting residues that had not been predicted by the initial CG simulations (Y^9.64^, Y^13.45^ and Y^13.51^ for *Tm*-PPase). However, these appear to only interact a proportionately small amount of the time compared to the lysine and arginine residues. In all these simulations there appeared to be no differences between the two interfacial binding sites in each mPPase or in the dynamics between the mutated and WT proteins throughout the atomistic simulations.

## Discussion

In this study, we have created atomistic models of three different mPPases in modelled membranes. Our work with the Vr-PPase created the first realistic model of Vr-PPase in a complex bilayer that mimics its native environment. Our simulations have shown that PPases inserted into a bilayer create an anionic fingerprint around them with preferential interactions with anionic lipids in two distinct sites. In mPPases, two pertinent studies of catalytic asymmetry and inter-subunit communication reported conflicting changes in PP_i_ affinity at the second monomer following binding of substrate at the first. One of the major differences in these experiments was whether the mPPases were in a lipid (Artukka et al., 2018) or detergent (Vidilaseris et al., 2019) environment. These data suggest that lipid binding could have a modulatory role in inter-subunit communication and catalytic asymmetry, and delipidation through solubilisation and purification procedures thereby influences mPPase function.

In this investigation, mPPases were found to interact preferentially with anionic lipids compared to neutral lipids. This interaction appears to be a result of the charged surface of the protein, as demonstrated by the electrostatic profile analysis and loss of interaction following mutation and loss of charge. In addition to this, the pronounced preference for POPA in *Tm*-PPase is likely due to its similarities with the *T. maritima* native DEGP lipids, as both are anionic and do not possess headgroups. This may also be the case for the *Cp*-PPase preferential interactions with POPA. Non-specific lipid interactions have been proposed to affect protein function. For example, cardiolipin binding to UraA (H^+^-uracil symporter) (Kalli et al., 2015), cytochrome *bc*_1_ (respiratory chain complex) (Arnarez et al., 2013) and SecY (proton translocon) (Corey et al., 2018) is thought to attract positive ions for pumping. It is unclear whether this functionality exists for other anionic lipids, as it is believed that this proton trap capability comes from the presence of two phosphate groups with different p*K*_a_ values (Kates et al., 1993). However, more recent studies suggest the p*K*_a_ values are similar for each cardiolipin phosphate (Olofsson and Sparr, 2013; Sathappa and Alder, 2016), so potentially this functionality is applicable to the single-phosphate anionic lipids in this investigation. Therefore, the anionic lipid clustering around the mPPases could serve to attract the pumped cations and the Mg_2_PP_i_ for catalysis.

The observed anionic interactions were localised to the cytoplasmic leaflet. These charged regions were shown to have a role in membrane protein insertion into the bilayer (Elazar et al., 2016). However, the clustering of anionic lipids to these areas and the specific interactions identified in this study may also play role in the function and stability of mPPases. Similar interactions between the lysine and arginine residues of the identified binding sites and the anionic lipid headgroups were also found in other proteins (Haviv et al., 2007; Vanegas and Arroyo, 2014; Norimatsu et al., 2017), in which the lipid interactions had marked effect on protein stability, function or dynamics. Additionally, it was more recently demonstrated for several oligomeric membrane-integral transporters that interfacial lipids play a crucial role in oligomerisation and stability (Gupta et al., 2017; Pyle et al., 2018). Lipids are also considered capable of stabilising transient conformational states (Sweadner, 2017). The need to stabilise alternative states of the catalytic cycle and members of the mPPase families means that MD-identified lipid interactions may be a key part of achieving structural information, particularly as interfacial binding sites were identified in this study.

Our studies identified two lipid binding sites on mPPases, an interfacial binding site and a distal binding site. The residues at the interfacial study are not highly conserved, ranging from 47.8% – 61.8% conservation. This is low for functionally relevant residues, but there are the repeated interactions with residues around K^9.7X^, such as K^9.70^ (*Tm*-PPase) and K^9.73^ (*Cp*-PPase), around K/R^13.52^ (K in *Tm*-PPase and *Vr*-PPase (K^13.48^) and R in *Cp*-PPase) and residues around K/R^14.4X^ (K^14.45^ in *Tm*-PPase, K^14.48^ in *Vr*-PPase and R^14.48^ in *Cp*-PPase). These differ in position by one helix turn, and so face out into the membrane, thereby preserving their function and potentially accommodating the change in headgroup size between POPA and PIP_2_. In addition to conservation of interactions with specific residues, they are positioned on functionally relevant helices. This clustering of interactions on TMH that form the dimer interface and are linked to inter-subunit communication and K^+^-dependency clearly supports our hypothesis that presence of anionic lipids at the interfacial interaction site may be functionally relevant to the mPPase catalytic cycle.

In addition to the computational evidence laid out in this work, there is experimental evidence of binding at the interfacial site. In the highest resolution mPPase structure currently available (*Vr*-PPase (PDB: 4A01)) (Lin et al., 2012), n-decyl-β-D-maltopyranoside (DM) detergent molecules are situated in this proposed binding site. In particular, the binding of one of these is highly similar to the anionic lipid positioning seen in our CG and AT simulations. Moreover, hydrogen bonds form between this detergent molecule and the K^9.70^ and K^10.49^ residues, mirroring the *Tm*-PPase simulations. The structural evidence of binding at the interfacial site in a mPPase provides further support to this region acting as a lipid binding site. The replacement of lipid with detergent was likely due to the purification and crystallisation conditions promoting removal of even tightly bound lipids (Yeagle, 2014). However, the binding of this detergent may have acted similarly to a lipid and helped maintain oligomeric stability, particularly as it has been suggested that detergents can bind in lipid interaction sites (Yeagle, 2014).

The interacting residues at the distal interaction site are not highly conserved (20.8% – 64%), but interactions cluster at similar structural positions. In all proteins in this study, interactions between anionic lipids and the proteins were formed with TMH 1, 2, 3 and 4, with R/K^1.60^ (R in *Tm*-PPase, and K in *Vr*-PPase *Cp*-PPase). Positively charged residues within one helix turn from R/K^1.60^, at K^1.61^ (*Tm*-PPase), K^1.62^ (*Vr*-PPase) and R^1.56^ (*Cp*-PPase) and residues in the midpoints of the helices: K^3.59^, R^3.63^ and R^4.48^ (*Tm*-PPase), K^2.50^, K^3.59^ and K^4.44^ (*Vr*-PPase), and K^2.46^, R^2.47^ and K^3.59^ (*Cp*-PPase) also formed large number of interactions with anionic lipids. In previous simulations of *Tm*-PPase, the distal region of the protein has been found to be flexible (Shah et al., 2017), and does not have high conservation between species. Therefore, our simulations suggest that the interactions are conserved to specific helix turns rather than individual residues. This might explain why despite the lower sequence similarity, we observe similar interactions with anionic lipids in the distal area. The distal lipid interaction sites may play a role in mPPase stability, as the region was identified during a case study of the IMPROvER (integral membrane protein stability selector) pipeline for stabilising mutations (Harborne et al., 2020). Further, three of the *Cp*-PPase residues identified by IMPROvER were found to interact with anionic lipids in our study (F^1.53^, R^3.67^ and R^7.62^). Due to the corroborative results between these two studies, the distal lipid interaction sites found in this work may play a role in protein stability.

Our study also provides evidence that lipid interactions in the distal and interfacial region may be a more general property of mPPase subfamilies, as *Tm*-PPase is a K^+^-dependent Na^+^-PPase, *Vr*-PPase is a K^+^-dependent H^+^-PPase and *Cp*-PPase is a K^+^-dependent Na^+^/H^+^-PPase. Despite the conserved pattern of interactions in our studies, there was no conserved sequence motif to identify these interactions in other family members, so likely homology modelling and electrostatic profile analysis, as performed here for *Cp*-PPase, would be required to identify binding sites in other homologues. The retention of interactions between pumping specificities is perhaps not unexpected, as the residues involved in coordinating the pumped ion are at the centre of the mPPase structure (Holmes et al., 2019). However, the conservation across subfamilies of lipid interactions that may stabilise the protein and be of mechanistic relevance bodes well for future structural and functional research using alternative homologues.

The preferential interaction of *Vr*-PPase with PIP_2_ in this work was striking, as PIP lipids are known to have roles in signalling, protein-protein interactions and are often found bound to proteins of interest (Corradi et al., 2018). The function of *Vr*-PPase in the tonoplast membrane of plants has been linked to auxin regulation and signalling (Li et al., 2005) and cooperative function with soluble pyrophosphatases (Segami et al., 2018); this, taken with the evidence of PIP_2_ binding in this study, may suggest a mechanism by which this signalling is able to take place. Additionally, the roles of the vacuolar ATPase complex and *Vr*-PPase are closely aligned (Holmes et al., 2019), which may also be mediated by PIP_2_ binding and activity.

This work provides the first evidence that interactions can form between mPPases and anionic lipids and that these are in regions that may hold functional significance and are conserved across homologues. These observations are very promising for future mPPase research. The role of the distal interaction site in stability needs to be investigated further, but co-crystallisation with lipids or mutagenesis to stabilise this region in detergent may be promising for structural studies and characterisation of other mPPases. Additionally, the putative role of the interfacial site in inter-subunit communication and catalysis warrants further investigation as it may be another factor in the apparent increasing complexity of the mPPase catalytic cycle.

## Methods

### Structure preparation and modelling

The coordinates of the resting-state crystal structures of *Tm*-PPase at 2.6 Å (PDB: 4AV3) (Kellosalo et al., 2012) and *Vr*-PPase at 3.5 Å (PDB: 5GPJ) (Li et al., 2016) were prepared for simulation by the addition of unresolved regions (residues 1, 30, 211-221, and 577-595 in *Tm*-PPase and 1-3, 39-62, and 262-278 in *Vr*-PPase) or mutations using Modeller (Webb and Sali, 2016). The structure of the mPPase from *Clostridium leptum* (*Cp*-PPase) was predicted by submitting the amino acid sequence to the Robetta (robetta.bakerlab.org) server for homology modelling (Song et al., 2013). The Ginzu domain prediction and structural homologue identification processes identified the 3.5 Å crystal structure of *Vr*-PPase (PDB: 5GPJ) (Li et al., 2016) as the most appropriate template for model building. Model quality analysis was performed by submitting the models to SWISS-MODEL Tools (swissmodel.expasy.org/qmean) for QMEAN (qualitive model energy analysis) and Z-score analysis using the QMEANBrane option (Studer et al., 2014). Modelling of *Cp*-PPase resulted in a similar structure.

### Coarse-grained molecular dynamics (CG-MD) simulations

All CG-MD simulations used the MARTINI 2.2 forcefield (Monticelli et al., 2008) and GROMACS 5.0.X (www.gromacs.org) (Van Der Spoel et al., 2005). The crystal structures or homology model were converted into coarse-grained (CG) models and centred independently in 16 x 16.5 x 16 nm^3^ simulation boxes. An elastic network was applied to the backbone atoms within 0.7 nm with a force constraint of 1000 kJmol^-1^nm^2^ to maintain the secondary and tertiary structure of the proteins. The CG proteins were energy minimised using the steepest descent algorithm embedded in GROMACS. Lipid bilayers were assembled around the proteins and the simulation boxes solvated by random placement of water and NaCl (150 mM) using insane protocols (Wassenaar et al., 2015). For *Tm*-PPase and *Cp*-PPase, the bilayers were made up of the palmitoyl-oleoyl phospholipid POPE in combination with anionic POPA, POPG or POPS. *Vr*-PPase was simulated in a bilayer composed of its native lipid constituents (cholesterol (29%), POPC (25%), POPE (17%), ceramide hexoside (17%), PIP_2_ (6%), POPG (3%), POPS (2%), POPA (1%)) (Yoshida and Uemura, 1986).

All systems were generated through this method as shown in Table 1. The complete lipidated and solvated systems were again energy minimised and equilibrated for 2 ns at 323 K, where the backbone particles were restrained. 5 independent repeat production simulations with different initial starting velocities were performed for 5 μs each. The temperature and pressure of the systems were maintained by a V-rescale thermostat (323 K) and a Parrinello-Rahman barostat (1 bar). The integration timestep was 20 fs.

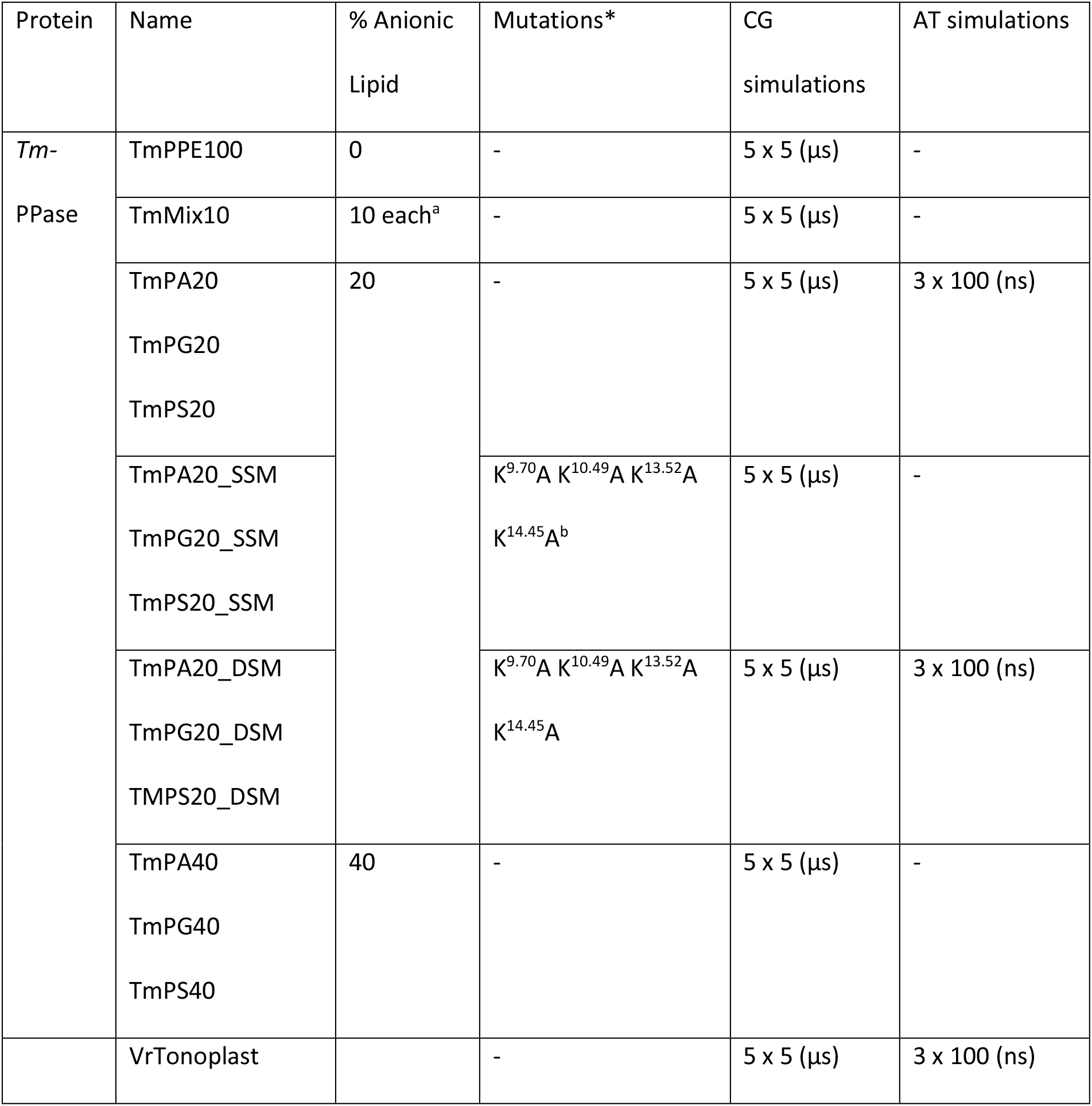
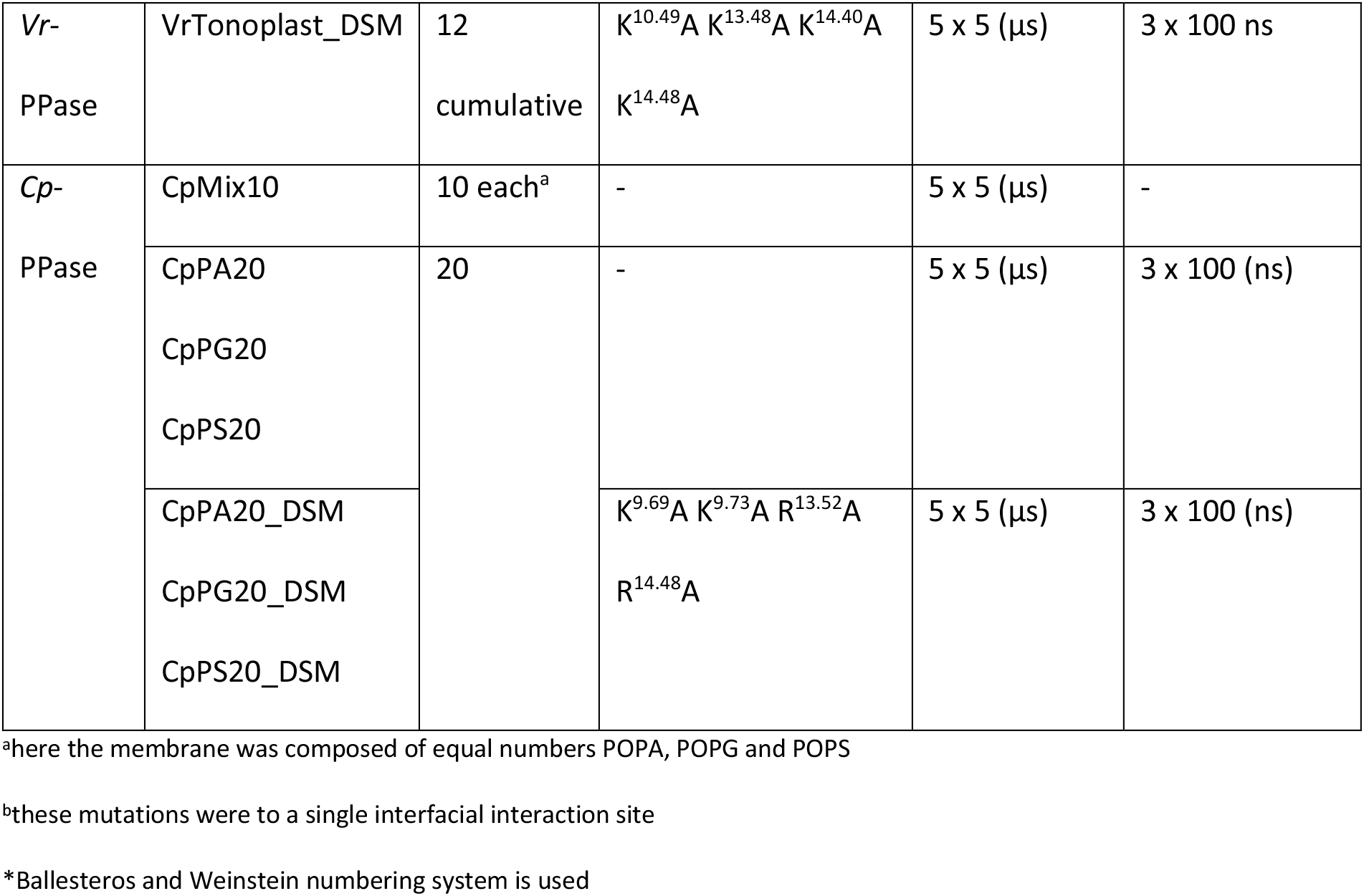

### CG-AT conversion

A serial multiscale approach was used, in which representative final frames of the selected CG simulations were converted to atomistic (AT) resolutions (“backmapping”). This conversion took place using the Backward protocol as described previously (Wassenaar et al., 2014). From each backmapped system, three replicates were generated with randomised starting velocities. These AT simulations were carried out using the CHARMM-36 forcefield (Brooks et al., 2009), a Nosé-Hoover thermostat (323 K), Parrenello-Rahman barostat (1 bar) and 2 fs timestep.

### Analysis

All analyses were performed using GROMACS (gmx mindist, gmx densmap, gmx rms, gmx rmsf), VMD (electrostatic profile, alignments) (Humphrey et al., 1996), PyMol (alignments and electron density inspection) (DeLano, 2002) and locally written scripts. For the contacts between lipid and protein, a locally written script calculated the interactions between each residue and the lipid head groups. A cut-off of 5.5 Å or 4 Å was defined for CG or AT simulations, respectively. For normalisation, all replicates were concatenated, and the number of contacts normalised to the total number of frames and number of that lipid species in the bilayer. The electrostatic profiles of protein structures were calculated by preparing the structure *via* PDB2PQR (Dolinsky et al., 2004) and then processed using the APBS plugin for VMD (Baker et al., 2001).

## Acknowledgements

This work was funded by a BBSRC DTP fellowship (BB/M011151/1) (to AH). AG was funded by a BBSRC grant BB/T006048/1 and a grant from the Academy of Finland, grant number 1322609. The work was also supported by access to the Advanced Research Computing facilities at the University of Leeds and the High-Performance Computing facilities from CSC, Finland.

## Supplementary Figures

**Supplementary Fig. 1.**
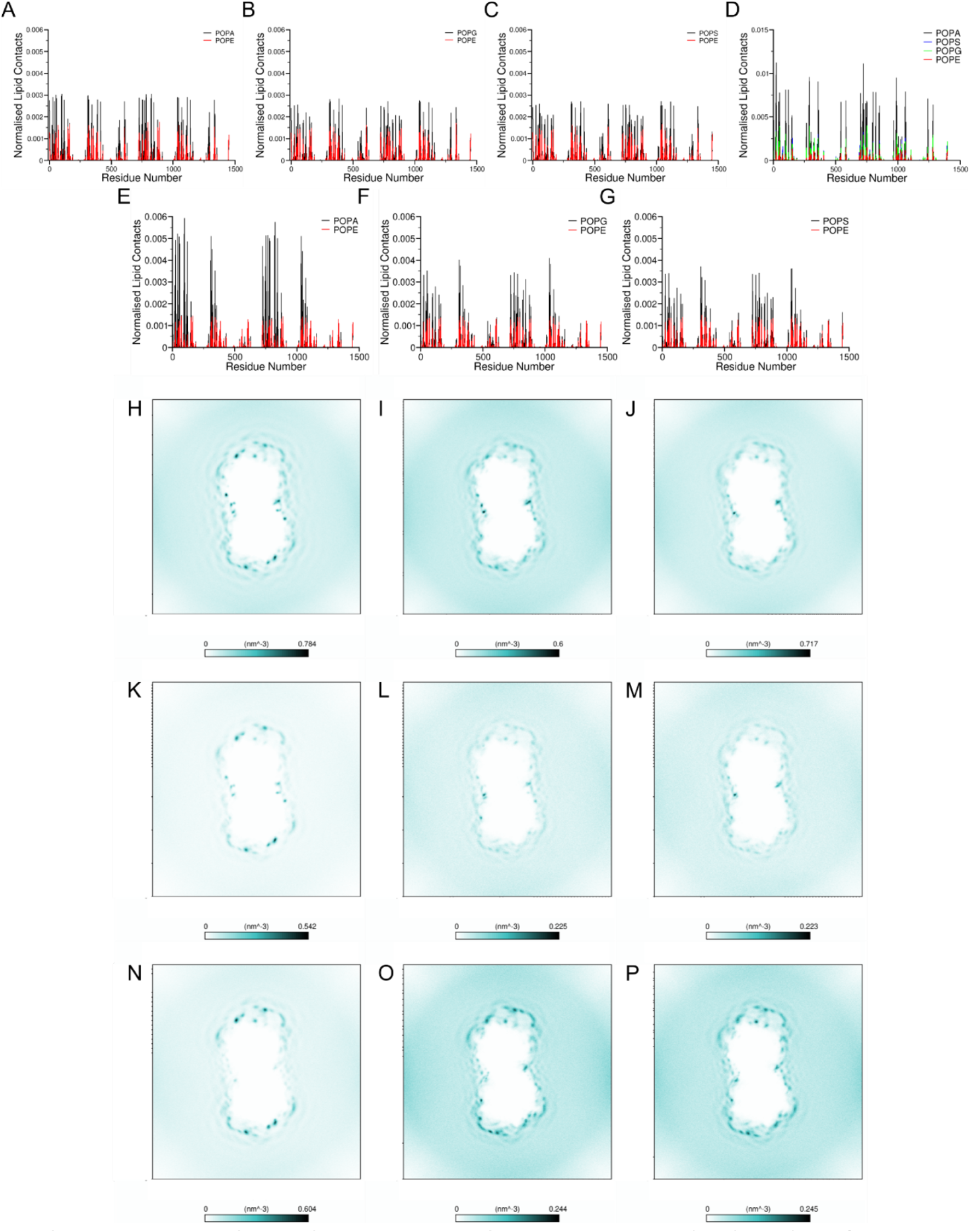
The Lipid Interactions with *Tm*-PPase. Normalised number of contacts between the lipids and *Tm*-PPase in the coarse-grained systems **A)** TmPA40, **B)** TmPG40, **C)** TmPS40, **D)** TmMix10, **E)** TmPA20_DSM **F)** TmPG20_DSM **G)** TmPS20_DSM. Density maps depicting the average density of phosphate particles of the anionic lipids in the **H)** TmPA40 **I)** TmPG40, **J)** TmPS40, **K-M)** POPA, POPG and POPS, respectively, from the TmMix10 system, and **N)** TmPA20_DSM **O)** TmPG20_DSM **P)** TmPS20_DSM systems.

**Supplementary Fig. 2.**
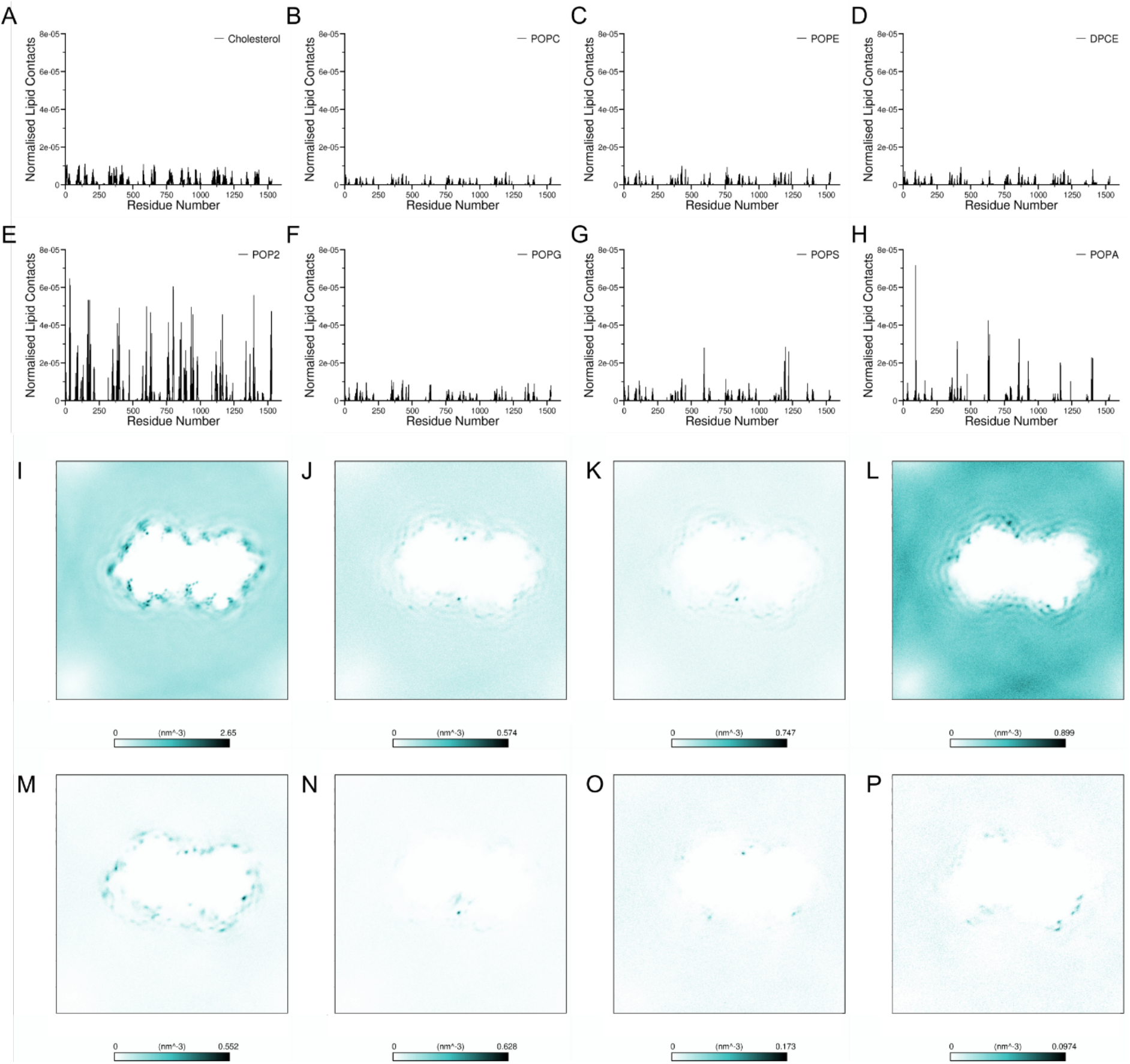
The Lipid Interactions with *Vr*-PPase. Normalised number of contacts between *Vr*-PPase during coarse-grained simulations and the bilayer representing a realistic tonoplast membrane comprised of **A)** Cholesterol, **B)** POPC, **C)** POPE, **D)** DPCE, **E)** POPG, **F)** POP2, **G)** POPS and **H)** POPA. Density maps depicting the average density of phosphate particles of the anionic lipids as in panels A-H.

**Supplementary Fig. 3.**
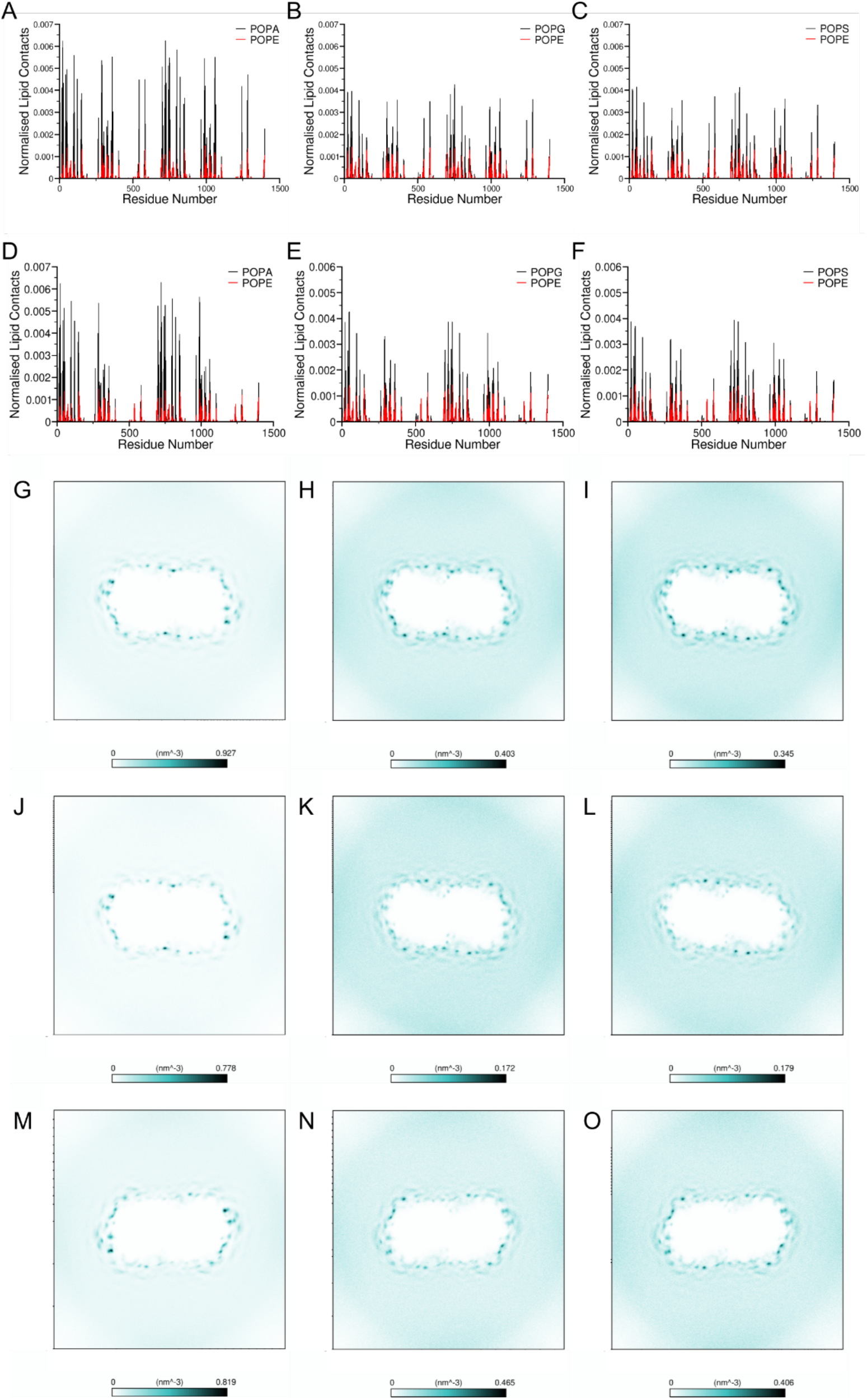
The Lipid Interactions with *Cp*-PPase. Normalised number of contacts between the lipids and *Cp*-PPase during coarse-grained simulations in the systems **A)** CpPA20, **B)** CpPG20, **C)** CpPS20, **D)** CpPA20_DSM, **E)** CpPG20_DSM and **F)** CpPS20_DSM. Density maps depicting the average density of phosphate particles of the anionic lipids in the systems **G)** CpPA20, **H)** CpPG20, **I)** CpPS20, **J-L)** POPA, POPG and POPS in CpMix10, **M)** CpPA20_DSM, **N)** CpPG20_DSM and **O)** CpPS20_DSM.

**Supplementary Fig. 4.**
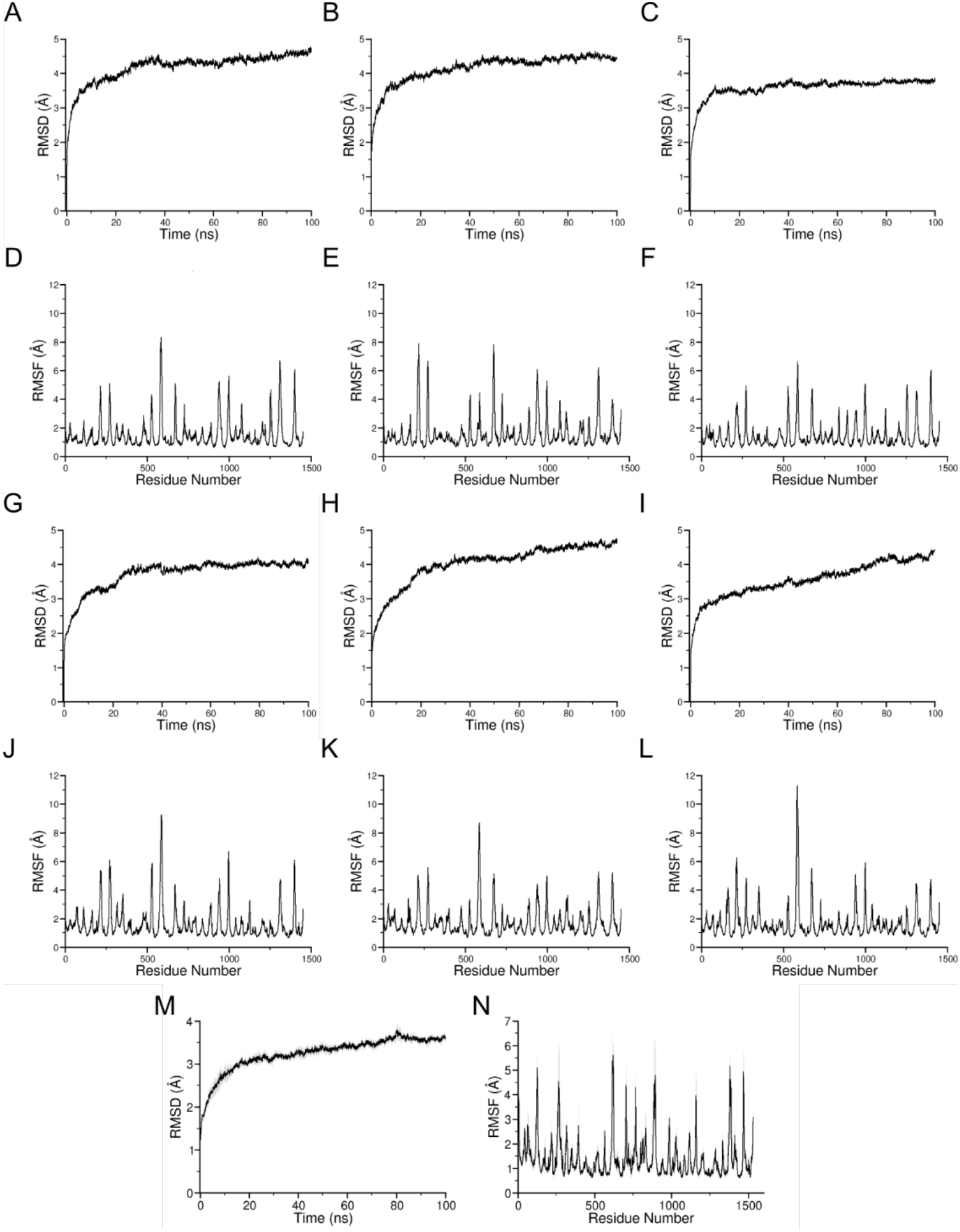
The Dynamics of Atomistic *Tm*-PPase and *Vr*-PPase. The RMSD/C_α_ of *Tm*-PPase during 100 ns of atomistic resolution simulation in systems **A)** TmPA20, **B)** TmPG20 and **C)** TmPS20, and **D-F)** RMSF/C_α_ of *Tm*-PPase in the same systems. **G-L)** The RMSD/C_α_ and RMSFC_α_ of the corresponding double interfacial site mutated version of *Tm*-PPase. **M)** The RMSD/C_α_ and **N)** RMSF/C_α_ of *Vr*-PPase during 100 ns of atomistic simulation in the tonoplast bilayer model.

**Supplementary Fig. 5.**
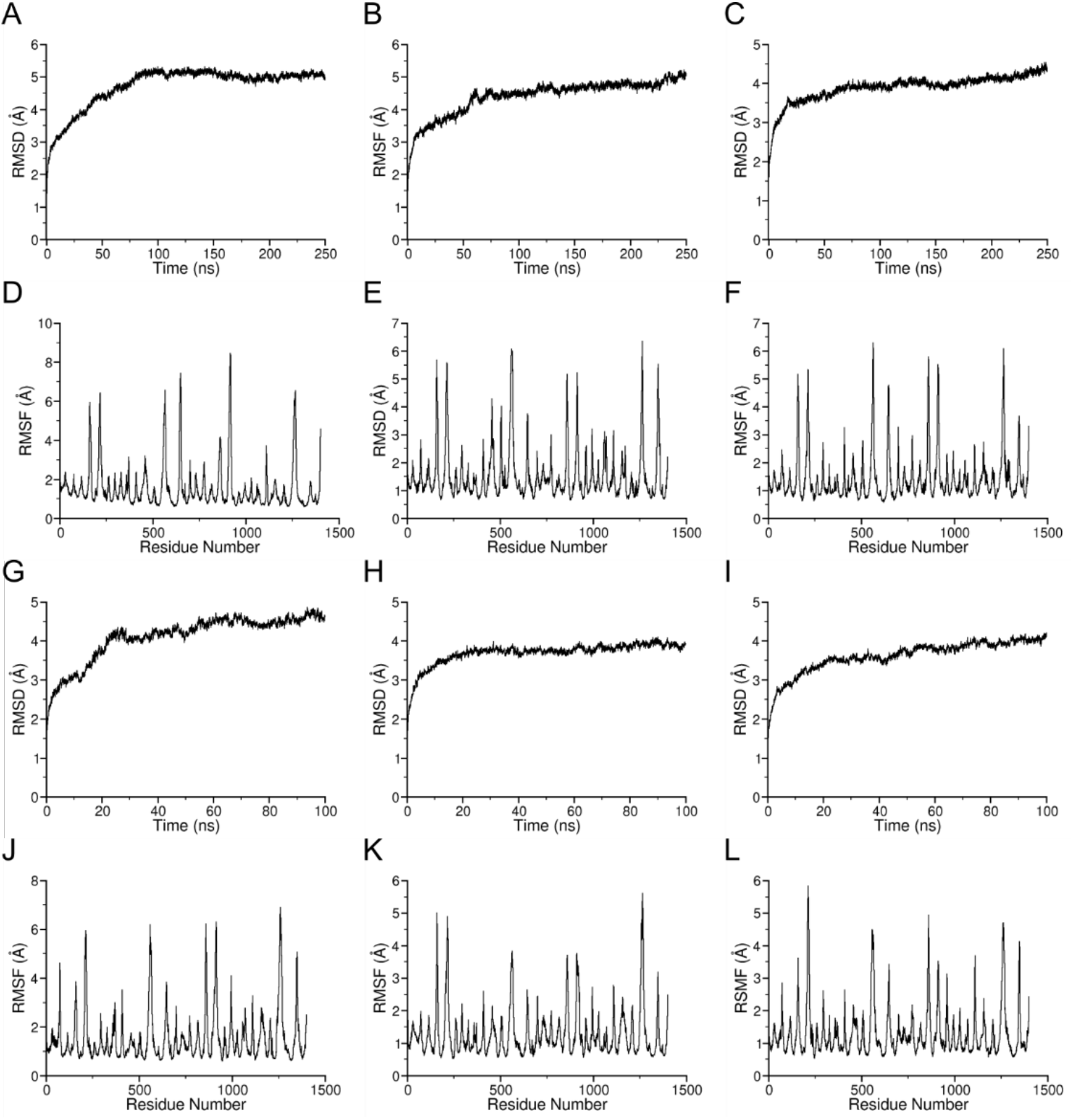
The Dynamics of Homology-Modelled Atomistic *Cp*-PPase. The RMSD/C_α_ of homology-modelled *Cp*-PPase during 100 ns of atomistic resolution simulation in systems **A)** CpPA20, **B)** CpPG20 and **C)** CpPS20, and **D-F)** RMSF/C_α_ of *Cp*-PPase in the same systems. **G-L)** The RMSD/C_α_ and RMSFC_α_ of the corresponding double interfacial site mutated version of *Cp*-PPase.

**Supplementary Fig. 6.**
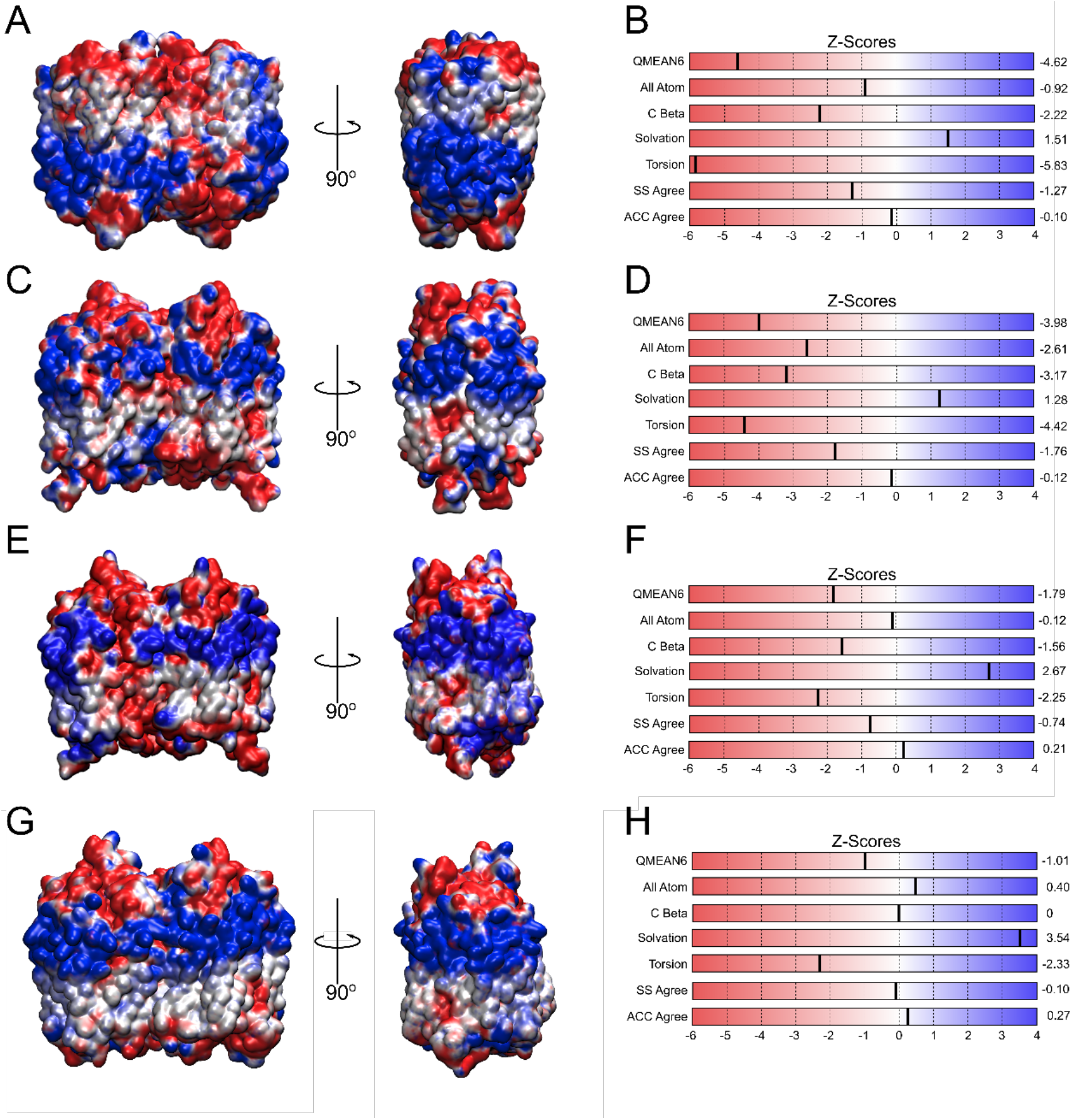
The Homology Models of *Cp*-PPase. The electrostatic profile and Z-Scores of the homology models of *Cp*-PPase generated by **A-B)** iTasser, **C-D)**, **E-F)** Robetta and **G-H)** AlphaFold2.

## Notes

### Competing Interest Statement

The authors have declared no competing interest.

## References

Anashkin, V.A., Malinen, A.M., Bogachev, A. V. and Baykov, A.A. 2021. Catalytic asymmetry in homodimeric H^+^-pumping membrane pyrophosphatase demonstrated by non-hydrolyzable pyrophosphate analogs. International Journal of Molecular Sciences. 22(18).

Arnarez, C., Mazat, J.-P., Elezgaray, J., Marrink, S.-J. and Periole, X. 2013. Evidence for Cardiolipin Binding Sites on the Membrane-Exposed Surface of the Cytochrome *bc*_1_. Journal of the American Chemical Society. 135(8), pp.3112–3120.

Artukka, E., Luoto, H.H., Baykov, A.A., Lahti, R. and Malinen, A.M. 2018. Role of the Potassium/Lysine Cationic Center in Catalysis and Functional Asymmetry in Membrane-Bound Pyrophosphatases. Biochemical Journal. 475(6), pp.1141–1158.

Au, K.M., Barabote, R.D., Hu, K.Y. and Jr, M.H.S. 2006. Evolutionary Appearance of H^+^-Translocating Pyrophosphatases. Microbiology. 152(5), pp.1243–1247.

Baker, N.A., Sept, D., Joseph, S., Holst, M.J. and McCammon, J.A. 2001. Electrostatics of nanosystems: Application to microtubules and the ribosome. Proceedings of the National Academy of Sciences of the United States of America. 98(18), pp.10037–10041.

Ballesteros, J.A. and Weinstein, H. 1995. Integrated Methods for the Construction of Three Dimensional Models and Computational Probing of Structure-Function Relations in G Protein-Coupled Receptors. Methods in Neurosciences. 25(19), pp.366–428.

Baykov, A.A. 2020. Energy Coupling in Cation-Pumping Pyrophosphatase—Back to Mitchell. Frontiers in Plant Science. 11(February), pp.1–5.

Bill, R.M., Henderson, P.J.F., Iwata, S., Kunji, E.R.S., Michel, H., Neutze, R., Newstead, S., Poolman, B., Tate, C.G. and Vogel, H. 2011. Overcoming Barriers to Membrane Protein Structure Determination. Nature Biotechnology. 29(4), pp.335–340.

Broecker, J., Eger, B.T. and Ernst, O.P. 2017. Crystallogenesis of Membrane Proteins Mediated by Polymer-Bounded Lipid Nanodiscs. Structure. 25(2), pp.384–392.

Brooks, B.R., Brooks, C.L., Mackerell, A.D., Nilsson, L., Petrella, R.J., Roux, B., Won, Y., Archontis, G., Bartels, C., Boresch, S., Caflisch, A., Caves, L., Cui, Q., Dinner, A.R., Feig, M., Fischer, S., Gao, J., Hodoscek, M., Im, W., Kuczera, K., Lazaridis, T., Ma, J., Ovchinnikov, V., Paci, E., Pastor, R.W., Post, C.B., Pu, J.Z., Schaefer, M., Tidor, B., Venable, R.M., Woodcock, H.L., Wu, X., Yang, W., York, D.M. and Karplus, M. 2009. CHARMM: The Biomolecular Simulation Program. Journal of Computational Chemistry. 30(10), pp.1545–1614.

Caffrey, M., Li, D. and Dukkipati, A. 2012. Membrane protein structure determination using crystallography and lipidic mesophases: Recent advances and successes. Biochemistry. 51(32), pp.6266–6288.

Cecchetti, C., Strauss, J., Stohrer, C., Naylor, C., Pryor, E., Hobbs, J., Tanley, S., Goldman, A. and Byrne, B. 2021. A novel high-throughput screen for identifying lipids that stabilise membrane proteins in detergent based solution. PLoS ONE. 16(7 July).

Corey, R.A., Pyle, E., Allen, W.J., Watkins, D.W., Casiraghi, M., Miroux, B., Arechaga, I., Politis, A. and Collinson, I. 2018. Specific Cardiolipin–SecY Interactions are Required for Proton-Motive Force Stimulation of Protein Secretion. Proceedings of the National Academy of Sciences. 115(31), pp.7967–7972.

Corradi, V., Mendez-Villuendas, E., Ingólfsson, H.I., Gu, R.X., Siuda, I., Melo, M.N., Moussatova, A., Degagné, L.J., Sejdiu, B.I., Singh, G., Wassenaar, T.A., Delgado Magnero, K., Marrink, S.J. and Tieleman, D.P. 2018. Lipid-Protein Interactions Are Unique Fingerprints for Membrane Proteins. ACS Central Science. 4(6), pp.709–717.

Corradi, V., Sejdiu, B.I., Mesa-Galloso, H., Abdizadeh, H., Noskov, S.Y., Marrink, S.J. and Tieleman, D.P. 2019. Emerging Diversity in Lipid-Protein Interactions. Chemical Reviews.

DeLano, W. 2002. Pymol: An Open-Source Molecular Graphics Tool. CCP4 Newsletter On Protein Crystallography. 40.

Dolinsky, T.J., Nielsen, J.E., McCammon, J.A. and Baker, N.A. 2004. PDB2PQR: An automated pipeline for the setup of Poisson-Boltzmann electrostatics calculations. Nucleic Acids Research. 32(WEB SERVER ISS.), pp.W665–W667.

Dörr, J.M., Scheidelaar, S., Koorengevel, M.C., Dominguez, J.J., Schäfer, M., van Walree, C.A. and Killian, J.A. 2016. The Styrene-Maleic Acid Copolymer: A Versatile Tool in Membrane Research. European Biophysics Journal. 45(1), pp.3–21.

Elazar, A., Weinstein, J.J., Prilusky, J. and Fleishman, S.J. 2016. Interplay Between Hydrophobicity and the Positive-Inside Rule in Determining Membrane-Protein Topology. Proceedings of the National Academy of Sciences. 113(37), pp.10340–10345.

Falkenburger, B.H., Jensen, J.B., Dickson, E.J., Suh, B.C. and Hille, B. 2010. Phosphoinositides: Lipid regulators of membrane proteins In: Journal of Physiology [Online]. J Physiol, pp.3179–3185. [Accessed 11 June 2021]. Available from: https://pubmed.ncbi.nlm.nih.gov/20519312/.

Guan, Z., Tian, B., Perfumo, A. and Goldfine, H. 2013. The polar lipids of Clostridium psychrophilum, an anaerobic psychrophile. Biochimica et Biophysica Acta - Molecular and Cell Biology of Lipids. 1831(6), pp.1108–1112.

Gupta, K., Donlan, J.A.C., Hopper, J.T.S., Uzdavinys, P., Landreh, M., Struwe, W.B., Drew, D., Baldwin, A.J., Stansfeld, P.J. and Robinson, C. V. 2017. The Role of Interfacial Lipids in Stabilizing Membrane Protein Oligomers. Nature. 541(7637), pp.421–424.

Habeck, M., Haviv, H., Katz, A., Kapri-Pardes, E., Ayciriex, S., Shevchenko, A., Ogawa, H., Toyoshima, C. and Karlish, S.J.D. 2015. Stimulation, Inhibition, or Stabilization of Na^+^,K^+^-ATPase Caused by Specific Lipid Interactions at Distinct Sites. Journal of Biological Chemistry. 290(8), pp.4829–4842.

Haines, T.H. and Dencher, N.A. 2002. Cardiolipin: A Proton Trap for Oxidative Phosphorylation. FEBS letters. 528(1–3), pp.35–39.

Harborne, S.P.D., Strauss, J., Boakes, J.C., Wright, D.L., Henderson, J.G., Boivineau, J., Jaakola, V.P. and Goldman, A. 2020. IMPROvER: the Integral Membrane Protein Stability Selector. Scientific Reports. 10(1), pp.1–18.

Haviv, H., Cohen, E., Lifshitz, Y., Tal, D.M., Goldshleger, R. and Karlish, S.J.D. 2007. Stabilization of Na^+^,K^+^-ATPase Purified from *Pichia pastoris* Membranes by Specific Interactions with Lipids. Biochemistry. 46(44), pp.12855–12867.

Haviv, H., Habeck, M., Kanai, R., Toyoshima, C. and Karlish, S.J.D. 2013. Neutral Phospholipids Stimulate Na^+^,K^+^-ATPase Activity: A Specific Lipid-Protein Interaction. Journal of Biological Chemistry. 288(14), pp.10073–10081.

Hedger, G., Koldsø, H., Chavent, M., Siebold, C., Rohatgi, R. and Sansom, M.S.P. 2019. Cholesterol Interaction Sites on the Transmembrane Domain of the Hedgehog Signal Transducer and Class F G Protein-Coupled Receptor Smoothened. Structure. 27(3), pp.549–559.e2.

von Heijne, G. 1992. Membrane Protein Structure Prediction. Hydrophobicity Analysis and the Positive-Inside Rule. Journal of molecular biology. 225(2), pp.487–94.

Holmes, A.O.M., Kalli, A.C., Goldman, A. and Goldman, A. 2019. The Function of Membrane Integral Pyrophosphatases From Whole Organism to Single Molecule. Frontiers in Molecular Biosciences. 6(November), pp.1–11.

Humphrey, W., Dalke, A. and Schulten, K. 1996. VMD: Visual Molecular Dynamics. Journal of Molecular Graphics. 14(1), pp.33–38.

Kajander, T., Kellosalo, J. and Goldman, A. 2013. Inorganic Pyrophosphatases: One Substrate, Three Mechanisms. FEBS Letters. 587(13), pp.1863–1869.

Kalli, A.C. and Reithmeier, R.A.F. 2018. Interaction of the human erythrocyte Band 3 anion exchanger 1 (AE1, SLC4A1) with lipids and glycophorin A: Molecular organization of the Wright (Wr) blood group antigen R. L. Dunbrack, ed. PLOS Computational Biology. 14(7), p.e1006284.

Kalli, A.C., Sansom, M.S.P. and Reithmeier, R.A.F. 2015. Molecular Dynamics Simulations of the Bacterial UraA H^+^-Uracil Symporter in Lipid Bilayers Reveal a Closed State and a Selective Interaction with Cardiolipin. PLoS Computational Biology. 11(3), pp.1–27.

Kates, M., Syz, J.Y., Gosser, D. and Haines, T.H. 1993. pH-Dissociation Characteristics of Cardiolipin and its 2’-Deoxy Analogue. Lipids. 28(10), pp.877–82.

Kellosalo, J., Kajander, T., Kogan, K., Pokharel, K. and Goldman, A. 2012. The Structure and Catalytic Cycle of a Sodium-Pumping Pyrophosphatase. Science. 337(6093), pp.473–476.

Lemercier, G., Dutoya, S., Luo, S., Ruiz, F.A., Rodrigues, C.O., Baltz, T., Docampo, R. and Bakalara, N. 2002. A Vacuolar-Type H^+^-Pyrophosphatase Governs Maintenance of Functional Acidocalcisomes and Growth of the Insect and Mammalian Forms of *Trypanosoma brucei*. Journal of Biological Chemistry. 277(40), pp.37369–37376.

Li, J., Yang, H., Peer, W.A., Richter, G., Titapiwantakun, B., Undurraga, S., Murphy, A.S., Gilroy, S. and Gaxiola, R. 2005. Arabidopsis H^+^-PPase AVP1 Regulates Auxin-Mediated Organ Development.. 310(5745), pp.121–125.

Li, K.M., Wilkinson, C., Kellosalo, J., Tsai, J.Y., Kajander, T., Jeuken, L.J.C., Sun, Y.J. and Goldman, A. 2016. Membrane Pyrophosphatases from *Thermotoga maritima* and *Vigna radiata* Suggest a Conserved Coupling Mechanism. Nature Communications. 7(13596), pp.1–11.

Lin, S.M., Tsai, J.Y., Hsiao, C.D., Huang, Y.T., Chiu, C.L., Liu, M.H., Tung, J.Y., Liu, T.H., Pan, R.L. and Sun, Y.J. 2012. Crystal Structure of a Membrane-Embedded H^+^-Translocating Pyrophosphatase. Nature. 484(7394), pp.399–403.

Liu, J., Pace, D., Dou, Z., King, T.P., Guidot, D., Li, Z.H., Carruthers, V.B. and Moreno, S.N.J. 2014. A Vacuolar-H^+^-Pyrophosphatase (TgVP1) is Required for Microneme Secretion, Host Cell Invasion, and Extracellular Survival of *Toxoplasma gondii*. Molecular Microbiology. 93(4), pp.698–712.

Luoto, H.H., Belogurov, G.A., Baykov, A.A., Lahti, R. and Malinen, A.M. 2011. Na^+^-Translocating Membrane Pyrophosphatases are Widespread in the Microbial World and Evolutionarily Precede H^+^-Translocating Pyrophosphatases. Journal of Biological Chemistry. 286(24), pp.21633–21642.

Luoto, H.H., Nordbo, E., Malinen, A.M., Baykov, A.A. and Lahti, R. 2015. Evolutionarily Divergent, Na^+^-Regulated H^+^-Transporting Membrane-Bound Pyrophosphatases. Biochemical Journal. 467(2), pp.281–291.

Monticelli, L., Kandasamy, S., Periole, X., Larson, R., Tieleman, D. and Marrink, S. 2008. {T}he {MARTINI} coarse{-}grained force field{:} {E}xtension to proteins. J. Chem. Theor. Comp. 4, pp.819–834.

Norimatsu, Y., Hasegawa, K., Shimizu, N. and Toyoshima, C. 2017. Protein-Phospholipid Interplay Revealed with Crystals of a Calcium Pump. Nature. 545(7653), pp.193–198.

Olofsson, G. and Sparr, E. 2013. Ionization Constants pKa of Cardiolipin M. Gasset, ed. PLoS ONE. 8(9), p.e73040.

Pollock, N.L., Lee, S.C., Patel, J.H., Gulamhussein, A.A. and Rothnie, A.J. 2018. Structure and Function of Membrane Proteins Encapsulated in a Polymer-Bound Lipid Bilayer. Biochimica et Biophysica Acta - Biomembranes. 1860(4), pp.809–817.

Pöyry, S. and Vattulainen, I. 2016. Role of charged lipids in membrane structures — Insight given by simulations. Biochimica et Biophysica Acta - Biomembranes. 1858(10), pp.2322–2333.

Pyle, E., Kalli, A.C., Amillis, S., Hall, Z., Lau, A.M., Hanyaloglu, A.C., Diallinas, G., Byrne, B. and Politis, A. 2018. Structural Lipids Enable the Formation of Functional Oligomers of the Eukaryotic Purine Symporter UapA. Cell Chemical Biology. 25(7), pp.840–848.

Rodrigues, C.O., Scott, D.A., Bailey, B.N., De Souza, W., Benchimol, M., Moreno, B., Urbina, J.A., Oldfield, E. and Moreno, S.N. 2000. Vacuolar Proton Pyrophosphatase Activity and Pyrophosphate (PP_i_) in *Toxoplasma gondii* as Possible Chemotherapeutic Targets. The Biochemical journal. 349(2000), pp.737–45.

Sathappa, M. and Alder, N.N. 2016. The Ionization Properties of Cardiolipin and its Variants in Model Bilayers. Biochimica et Biophysica Acta (BBA) - Biomembranes. 1858(6), pp.1362–1372.

Segami, S., Tomoyama, T., Sakamoto, S., Gunji, S., Fukuda, M., Kinoshita, S., Mitsuda, N., Ferjani, A. and Maeshima, M. 2018. Vacuolar H^+^-Pyrophosphatase and Cytosolic Soluble Pyrophosphatases Cooperatively Regulate Pyrophosphate Levels in *Arabidopsis thaliana*. The Plant Cell. 30(5), pp.1040–1061.

Shah, N.R., Wilkinson, C., Harborne, S.P.D., Turku, A., Li, K.M., Sun, Y.J., Harris, S. and Goldman, A. 2017. Insights into the Mechanism of Membrane Pyrophosphatases by Combining Experiment and Computer Simulation. Structural Dynamics. 4(3), pp.1–12.

Sohlenkamp, C. and Geiger, O. 2015. Bacterial membrane lipids: Diversity in structures and pathways. FEMS Microbiology Reviews. 40(1), pp.133–159.

Song, Y., Dimaio, F., Wang, R.Y.R., Kim, D., Miles, C., Brunette, T., Thompson, J. and Baker, D. 2013. High-resolution comparative modeling with RosettaCM. Structure. 21(10), pp.1735–1742.

Van Der Spoel, D., Lindahl, E., Hess, B., Groenhof, G., Mark, A.E. and Berendsen, H.J.C. 2005. GROMACS: Fast, Flexible, and Free. Journal of Computational Chemistry. 26(16), pp.1701–1718.

Stansfeld, P.J., Jefferys, E.E. and Sansom, M.S.P. 2013. Multiscale simulations reveal conserved patterns of lipid interactions with aquaporins. Structure. 21(5), pp.810–819.

Studer, G., Biasini, M. and Schwede, T. 2014. Assessing the local structural quality of transmembrane protein models using statistical potentials (QMEANBrane). Bioinformatics. 30(17), pp.i505–i511.

Sweadner, K.J. 2017. An Ion-Transport Enzyme That Rocks. Nature. 545(7653), pp.162–164.

Takeda, Y. and Kasamo, K. 2002. Transmembrane topography of plasma membrane constituents in mung bean (Vigna radiata L.) hypocotyl cells. II. The large scale asymmetry of surface peptides. Biochimica et Biophysica Acta - Biomembranes. 1558(1), pp.14–25.

Tsai, J.-Y., Tang, K.-Z., Li, K.-M., Hsu, B.-L., Chiang, Y.-W., Goldman, A. and Sun, Y.-J. 2019. Roles of the Hydrophobic Gate and Exit Channel in *Vigna radiata* Pyrophosphatase Ion Translocation. Journal of Molecular Biology. 431(8), pp.1619–1632.

Tsai, J.Y., Kellosalo, J., Sun, Y.J. and Goldman, A. 2014. Proton/Sodium Pumping Pyrophosphatases: The Last of the Primary Ion Pumps. Current Opinion in Structural Biology. 27(1), pp.38–47.

Vanegas, J.M. and Arroyo, M. 2014. Force Transduction and Lipid Binding in MscL: A Continuum-Molecular Approach D. Fotiadis, ed. PLoS ONE. 9(12), p.e113947.

Vidilaseris, K., Kiriazis, A., Turku, A., Khattab, A., Johansson, N.G., Leino, T.O., Kiuru, P.S., Boije af Gennäs, G., Meri, S., Yli-Kauhaluoma, J., Xhaard, H. and Goldman, A. 2019. Asymmetry in Catalysis by *Thermotoga maritima* Membrane-Bound Pyrophosphatase Demonstrated by a Nonphosphorus Allosteric Inhibitor. Science Advances. 5(5), p.eaav7574.

Völkl, P., Huber, R., Drobner, E., Rachel, R., Burggraf, S., Trincone, A. and Stetter, K.O. 1993. Pyrobaculum aerophilum sp. nov., a novel nitrate-reducing hyperthermophilic archaeum. Applied and Environmental Microbiology. 59(9).

Wassenaar, T.A., Pluhackova, K., Böckmann, R.A., Marrink, S.J. and Tieleman, D.P. 2014. Going Backward: A Flexible Geometric Approach to Reverse Transformation from Coarse Grained to Atomistic Models. Journal of Chemical Theory and Computation. 10(2), pp.676–690.

Wassenaar, T.A., Ingólfsson, H.I., Böckmann, R.A., Tieleman, D.P. and Marrink, S.J. 2015. Computational lipidomics with insane: A versatile tool for generating custom membranes for molecular simulations. Journal of Chemical Theory and Computation. 11(5), pp.2144–2155.

Webb, B. and Sali, A. 2016. Comparative Protein Structure Modeling Using MODELLER. Current Protocols in Bioinformatics. 54(1), pp.1–37.

Yeagle, P.L. 2014. Non-Covalent Binding of Membrane Lipids to Membrane Proteins. Biochimica et Biophysica Acta - Biomembranes. 1838(6), pp.1548–1559.

Yoon, H.S., Kim, S.Y. and Kim, I.S. 2013. Stress Response of Plant H^+^-PPase-Expressing Transgenic *Escherichia coli* and *Saccharomyces cerevisiae:* A Potentially Useful Mechanism for the Development of Stress-Tolerant Organisms. Journal of Applied Genetics. 54(1), pp.129–133.

Yoshida, S. and Uemura, M. 1986. Lipid Composition of Plasma Membranes and Tonoplasts Isolated from Etiolated Seedlings of Mung Bean (*Vigna radiata* L.). Plant Physiology. 82(3), pp.807–812.

